# Comparative insights to the transportome of *Nosema*: a genus of parasitic microsporidians

**DOI:** 10.1101/110809

**Authors:** Hasnahana Chetia, Debajyoti Kabiraj, Swagata Sharma, Utpal Bora

## Abstract

*Nosema,* a genus of parasitic microsporidia, causes pebrine disease in arthropods, including economically important silkworms and honeybees. *Nosema* have gene-poor genomes shaped by loss of the metabolic pathways, as a consequence of continued dependence on host-derived substrates. As an act of counterbalance, they have developed an array of transporter proteins that allow stealing from their hosts. Here, we have identified the core set of twelve transporter families present in *Nosema* genus, viz. *N. apis, N. bombycis, N. ceranae* and *N. antheraea* through *in silico* pipeline. Transportomes of *N. apis, N. bombycis, N. ceranae* and *N. antheraea* have a dominant share of secondary carriers and primary active transporters. The comparatively rich and diverse transportome of *N. bombycis* indicates the role of transporters in its remarkable capability of host adaptation. The core set of transporter families of *Nosema* includes ones that have a likely role in osmo-regulation, intra- and extra-cellular pH regulation, energy compensation and self-defence mechanism. This study has also revealed a set of ten species-specific transporter families within the genus. To our knowledge, this is the first ever intra-genus study on microsporidian transporters. Both these datasets constitutes a valuable resource that can aid in development of inhibitor-based *Nosema* management strategies.

## [1] Introduction

Microsporidia is a specific group of unicellular obligate parasites or hyperparasites which can infect a myriad of organisms including a few economically important insects like silkworms, honey bees etc. as well as humans **(Silveira *et al.,* 2009; Weiss and Becnel, 2014)**. They generally have highly reduced gene-dense genomes achieved via evolutionary sacrifice of the genes of many essential pathways (TCA cycle, metabolic pathways) to minimize biological complexity **(Keeling and Corradi, 2011; Katinka *et al.,* 2001)**. Microsporidia possess an unstacked Golgi apparatus and a cryptic genome-less mitochondria called mitosome. In order to compensate for their reduced metabolic capacity, they utilize the host’s inner metabolism to derive nutrition. They usually do this either by up-regulating the host’s metabolic pathways via microsporidia-secreted factors and taking up nutrients from the host by their membrane-bound transporters **(Cuomo *et al.,* 2012; Heinz *et al.,* 2014)**. The pathogenic abilities of microsporidia are extraordinary as they are laden with an armoury to invade and survive inside host cells. One important weapon in this armoury is the microsporidian transporters that span across membranes, facilitating exchange of substrates between host and microsporidia, thus helping in its sustenance the host cytoplasm. This way, it brilliantly bypasses the energy-expensive biochemical pathways. An example of microsporidian transporters are the ATP transporters ubiquitously present in all microsporidia known **(Corradi, 2015)**. Genomic analyses have shown that despite undergoing genome reduction, a set of core transporter proteins are retained by microsporidia which are similar to other parasites **(Heinz *et al.,* 2012)**. These core transporters make the influx of nicotinic acids, cations, sugars, ATP etc. possible within the body of the parasite to help it in survival and proliferation. It has also been found that transporter proteins are over-expressed in microsporidian genomes **(Peyretaillade *et al.,* 2012).** Though the role of microsporidian transporters in host specificity, adaptation and interaction is apparent, less is understood about how a host or its environment can shape the transportome of a microsporidia. Studying the whole transportome of a microsporidia can provide a head start to deciphering these concealed aspects.

The focus of present study was the genus *Nosema* (Nosematidae family) which consists of microsporidian species that generally infects insects. It is the most diverse group of microsporidia with 150 genera reported so far **(Wittner, 1999)**. The disease caused by *Nosema* is broadly called Nosemosis (after the genus) or Pebrine, where destruction of insect midgut epithelial cells is a common pathological manifestation **(Hges *et al.,* 2010)**. *Nosema* infect insects from different orders such as Hymenoptera (honey bee), Lepidoptera (silkworms), etc **(Chen and Huang, 2010)**. Huge economic losses are incurred by countries of Europe, Asia and others every year due to mass death of these economically important host organisms which are utilized in sericulture, apiculture etc **(Farrar, 1947; Jeffree and Allen, 1956)**. At present, the whole genome for three *Nosema* species, namely, *Nosema apis, N. bombycis* and *N. ceranae* is available in the NCBI Genome database **(Table-1)**. The first genome to be published was that of *N. ceranae* in 2009 with a draft assembly size of 7.86 Mb **(Cornman *et al.,* 2009) (Table-1)**. It has a highly AT-biased and genetically diverse genome, rich in repetitive elements **(Roudel *et al.,* 2013)**. *N. ceranae* has been found to be associated with “colony collapse disorder” in European honey bees, *Apis mellifera,* causing huge economic losses in apiculture sector **(Higes *et al.,* 2008)**. Since honey bees are principle pollinators, their mass deaths have also affected agricultural productivity and sustainability. The second *Nosema* species with a sequenced whole genome is *N. bombycis.* It has a GC-rich (~33%), gene-dense genome (~15.69 Mb) which is among the largest known microsporidian genomes **(Table-1)**. Its aforementioned gene density has been attributed by abundant gene duplications, host-derived transposomal element proliferation and acquisition of bacterial genes via horizontal gene transfer (HGT) **(Pan *et al.,* 2013)**. It is the causal organism of pebrine disease in domestic silkworms causing significant losses in sericulture industry **(Ishihara and Fujiwara, 1965)**. *N. apis* is another microsporidia responsible for death of European honey bees. The genome of *N. apis* was assembled via shotgun sequencing and its size is 8.5 Mbp **(Chen *et al.,* 2013) (Table-1)**. Though it is thought to be lower in virulence than *N. ceranae, N. apis* also plays a significant role in causing mass fatality among honey bee colonies. *N. ceranae* and *N. apis* are phylogenetically closer as compared to *N. bombycis* **(Chen and Huang, 2010)**. Apart from these three species, *N. antheraea* (also known as *N. pernyi)* which infects the wild and undomesticated tassar moth, *Antheraea pernyi,* is also studied as a causal organism of pebrine disease in silkworms (**Pan *et al*., 2013**) (**Table-1**).

**Table 1.**
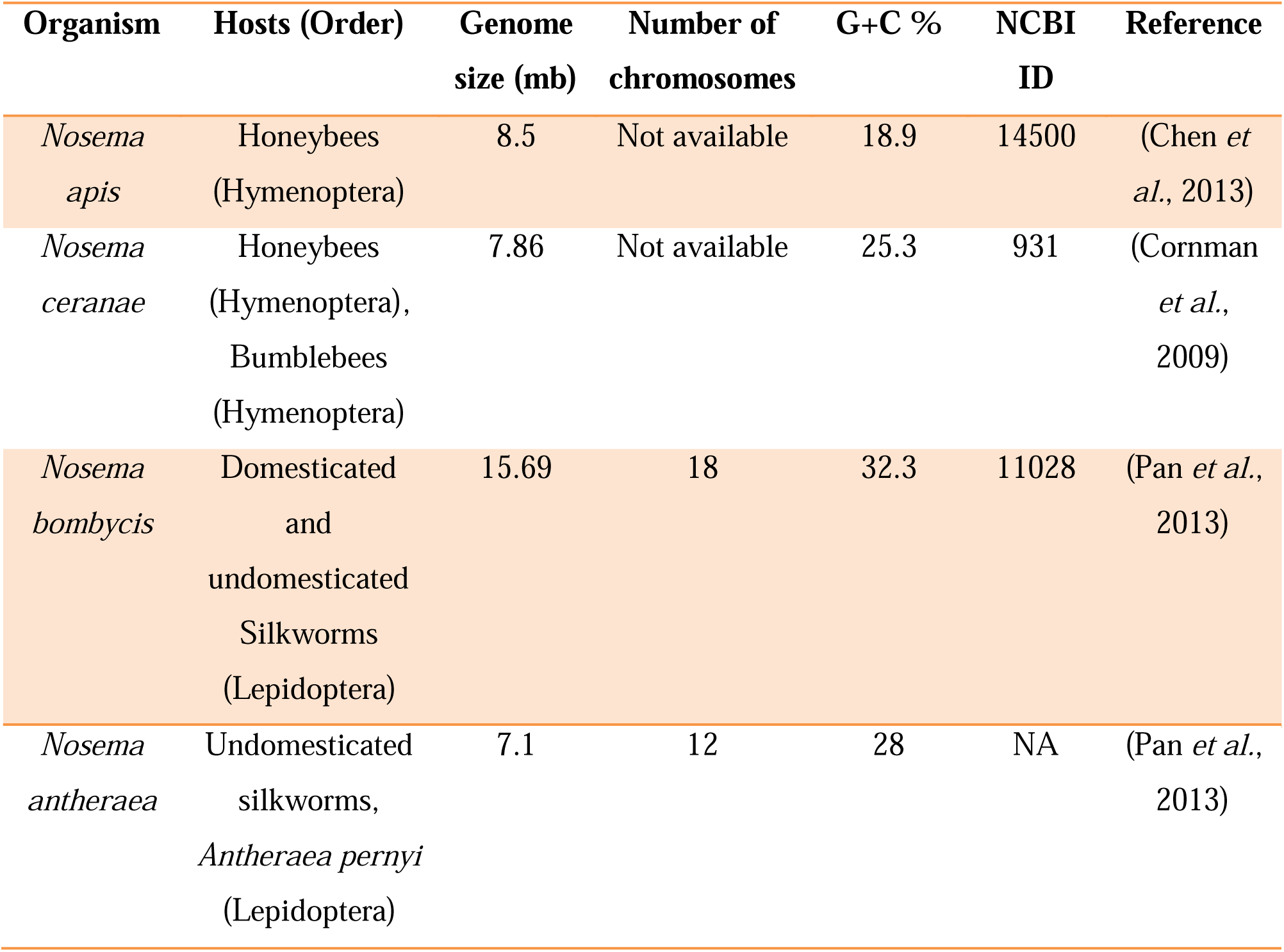
Comparative genomic information of *Nosema apis, N. ceranae, N. bombycis* and *N. antheraea*.

| Organism | Hosts (Order) | Genome size (mb) | Number of chromosomes | G+C % | NCBI ID | Reference |
| --- | --- | --- | --- | --- | --- | --- |
| *Nosema apis* | Honeybees (Hymenoptera) | 8.5 | Not available | 18.9 | 14500 | (Chen et al., 2013) |
| *Nosema ceranae* | Honeybees (Hymenoptera), Bumblebees (Hymenoptera) | 7.86 | Not available | 25.3 | 931 | (Cornman et al., 2009) |
| *Nosema bombycis* | Domesticated and undomesticated Silkworms (Lepidoptera) | 15.69 | 18 | 32.3 | 11028 | (Pan et al., 2013) |
| *Nosema* | Undomesticated | 7.1 | 12 | 28 | NA | (Pan *et al.*, |
| *antheraea* | silkworms, |  |  |  |  | 2013) |
|  | *Antheraea pernyi* |  |  |  |  |  |
|  | (Lepidoptera) |  |  |  |  |  |

In this study, we retrieved the available proteome data on the four species of *Nosema (N. apis, N. bombycis, N. ceranae* and *N. antheraea)* and identified the transporter proteins within the dataset using an *in silico* workflow. Furthermore, we classified the complete transportome of these four organisms into channels, secondary carriers and primary transporters based on the classification of TCDB (Transporter Classification Database) **(Saier *et al.,* 2006)**. Intra-genus comparison revealed a core set of *Nosema* transporters involved in transport of ATP, sugar, cations, etc. Our study provides a snapshot of the transportome of *Nosema* genus whose comparison to existing data on microsporidia provides a delineated view of the conserved and unique aspects of this genus.

## [2] Methodology

The protein sequences of *N. apis* BRL01, *N. bombycis* CQ1 and *N. ceranae* BRL01 were retrieved from UniProt database **(The UniProt Consortium, 2014)**. Similarly, the draft proteome of *N. antheraea* (Isolate YY, version 1) consisting of predicted ORFs (Open reading frames) was obtained from SilkPathDB **(Pan *et al.,* 2013)**. Using TCDB’s ftp link, its mapped dataset of transporter proteins was downloaded. A local standalone BlastP program was run using the TCDB dataset as a “blastdb” (database) and the four proteomes as query with an e-value cut off 0.001 **(Saier *et al.,* 2006)**. The proteins with a percentage similarity of less than 30% were discarded. Then, the proteins were further filtered out on the basis of presence or absence of transmembrane (TM) domains. The TM domains were detected using three different web-servers specifically used for TM prediction, namely, HMMTOP, TMHMMv2.0 and Phobius with default parameters **(Krogh *et al.,* 2001; Tusnády and Simon, 2001; Käll *et al.,* 2007)**. Only those proteins whose TM domains were predicted by at least two of the three tools were selected for further analysis. Furthermore, a two-way analysis of conserved domains of the filtered proteins was carried out using a local installation of InterProScan (version 5.17) and the web-server CD-Search based on CDD (Conserved Domain Database) **(Jones *et al.,* 2014; Marchler-Bauer *et al.,* 2014)**. Only those proteins whose predicted domains from TCDB families matched with the predicted domains of InterProScan and CDD were retained as putative, *in silico* characterized transporters. These transporters were classified according the TCDB nomenclature (till the third position to indicate its respective family or superfamily). The flow chart representing the method and online/offline tools used in this study is shown in **Figure 1**.

**Figure 1.**
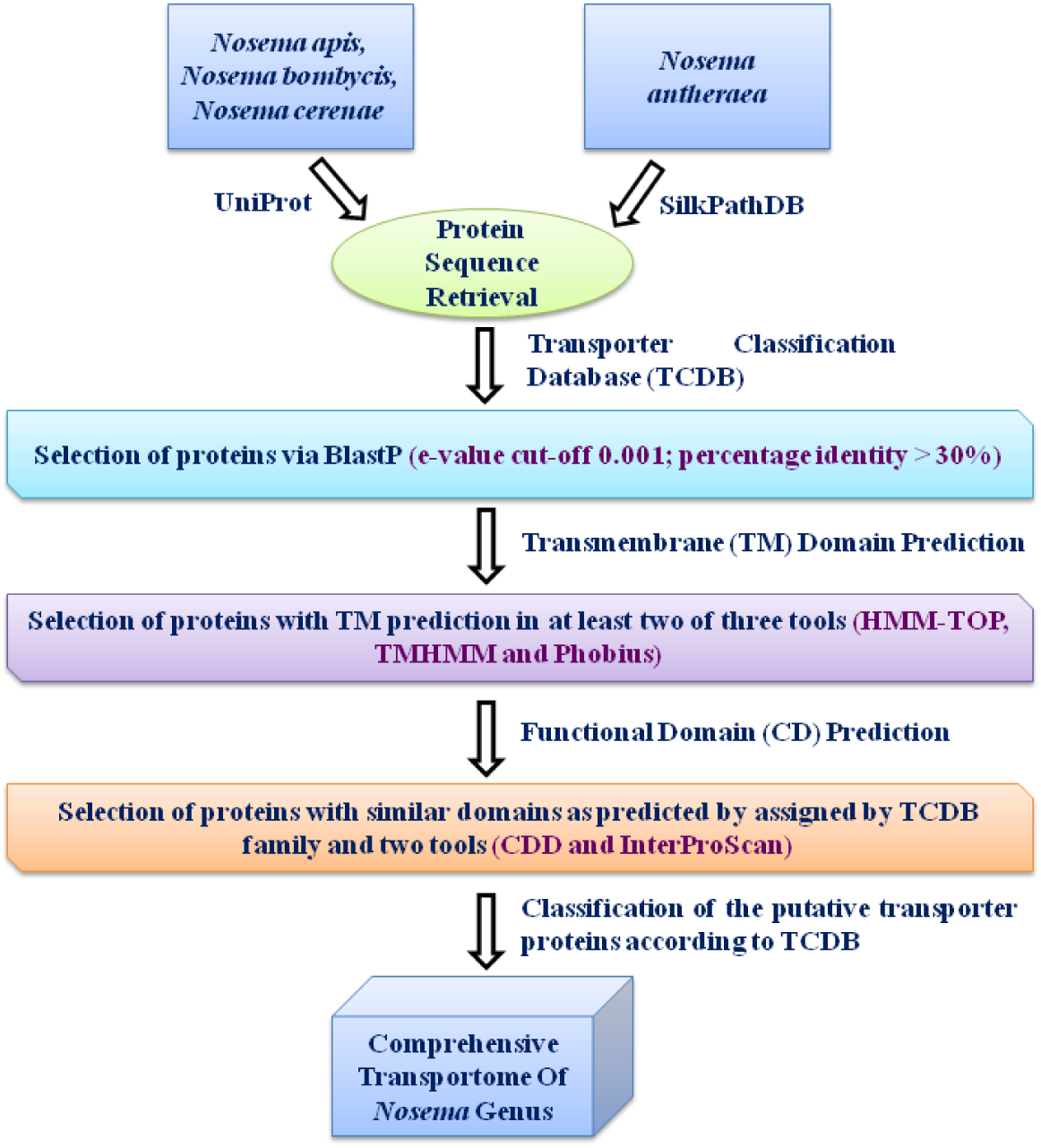
Workflow used for identification and classification of the complete transportome of *Nosema* genus.

## [3] Results and Discussion

Our analysis showed that the transportome of *N. apis, N. bombycis, N. ceranae* and *N. antheraea* consisted of 41, 78, 47 and 51 proteins respectively **(Refer to S-1)**. Secondary carrier type facilitators (Class 2) were found to be the most abundant class of transporters in *N. apis, N. bombycis* and *N. ceranae* while primary transporters (Class 1) was found to be most abundant in *N. antheraea* **(Figure 2)**. Class 8 and 9 contains accessory proteins to other transporters and uncharacterized transporters, respectively, so they were not probed further in this study. The distribution of the transporter protein families and their substrates in *N. apis, N. bombycis, N. ceranae* and *N. antheraea* has been shown in **Table 2**, **3** and **4**.

**Figure 2.**
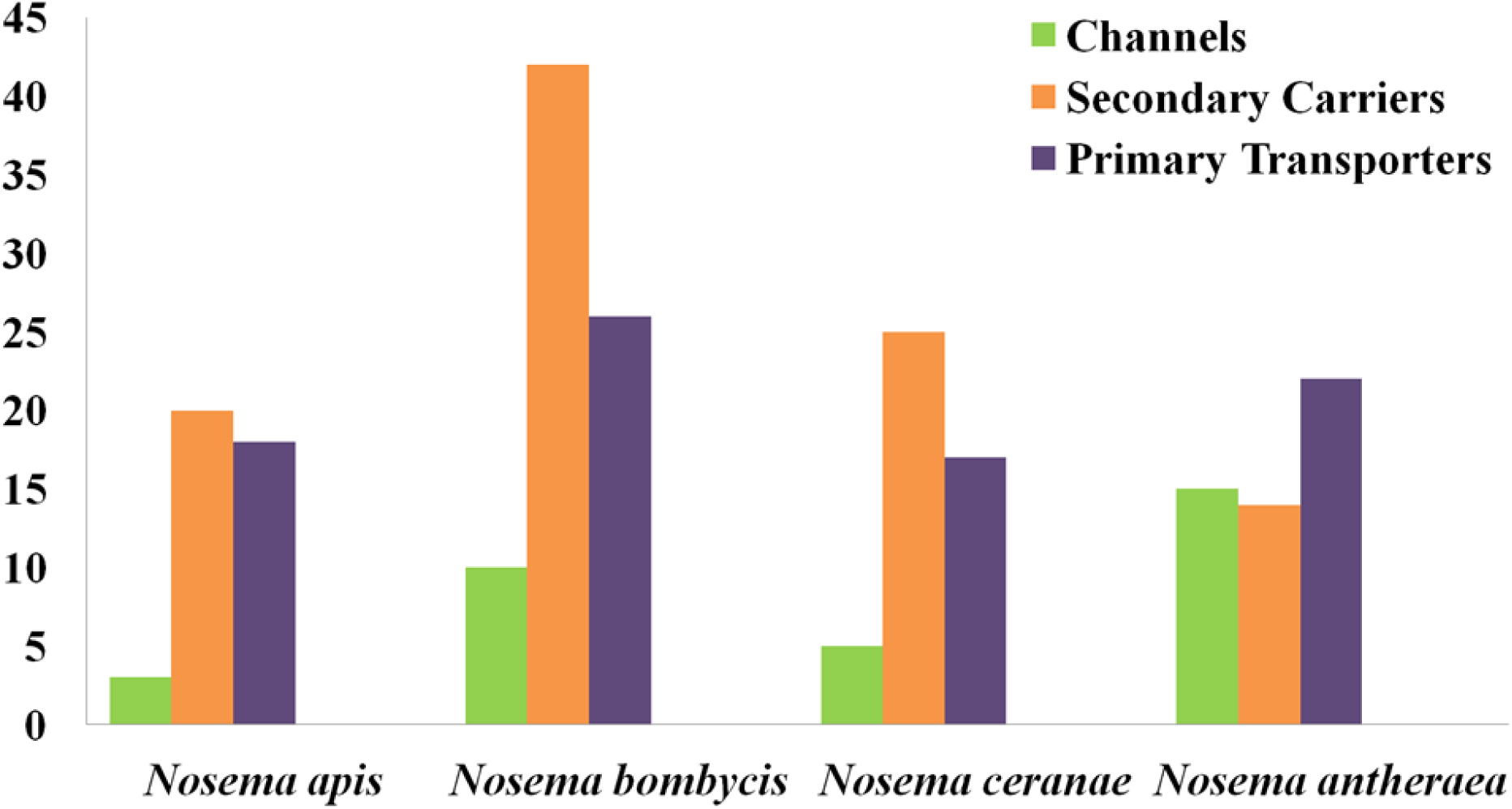
Class-wise distribution of the transporter proteins constituting the transportome of *Nosema apis, N. bombycis, N. ceranae* and *N. antheraea*.

**Table 2.**
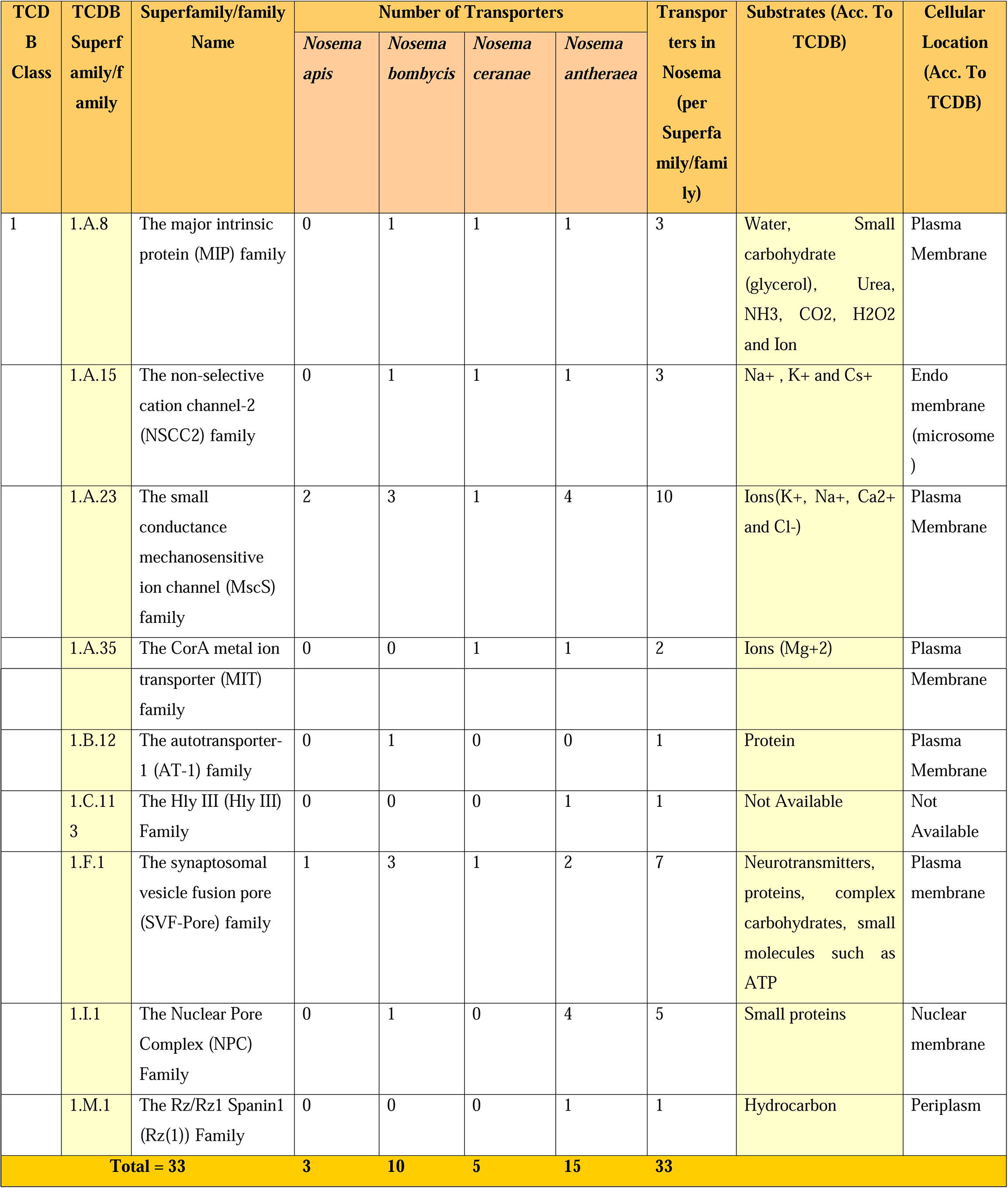
Distribution of Class 1 transporters with their substrates for *Nosema apis, N. bombycis*, *N. ceranae* and *N. antheraea:*

### [3.1] Class 1: Channels and Pores

Channels or pores which constitute the Class-1 of TCDB facilitate passive transport of substrates without the involvement of additional reaction and represents passive transport. Class 1 has transporting channels mostly made up of α-helices or β-sheets spanning the lipid bilayer and forming a channel or pore in the organism’s cell membrane to allow passage of solutes etc **(Saier *et al.,* 2006)**. *Nosema* being a highly reduced obligate parasite lacks many basic biosynthetic pathways and relies upon its host for requirements like nucleotides which are building blocks of nucleic acid, etc **(Heinz *et al.,* 2014)**. Thus, Class 1 transporters, which don’t require ATP, should be a preferred choice for this organism. Our study showed that a total of thirty-three proteins from the four *Nosema* species belonged to Class 1 out of which α-type channels were the most commonly expressed transporters **(Table 2)**. *N. antheraea* had the largest arsenal of channels (15 no.s) among the four species and *N. apis* had the least (3 no.s).

All four *Nosema* sp.s uniformly expressed transporters from classes named “Small Conductance Mechanosensitive Ion Channel (MscS) Family” (1.A.23) and “Synaptosomal Vesicle Fusion Pore (SVF-Pore) Family” (1.F.1). Previous studies have revealed that MscS transporters are conserved within other microsporidia such as *E. cuniculi, E. bieneusi* and *A. algerae* **(Peyretaillade *et al.,* 2012)**. The MscS transporters represent a modest subset of *Nosema* transporters which are responsive to mechanical perturbations in the lipid bilayer, acting as mechanical switches. Their conservation within four *Nosema* genomes indicates that these proteins can be highly helpful in acquiring surface-located or secreted host proteins **(Nakjang *et al.,* 2013)**. It also suggests that these proteins might be essential for fulfilling the core features of a parasite life cycle such as general mechanisms of host exploitation. Furthermore, the SVF-Pore family (1.F.1), which was discovered uniformly in all four *Nosema* species, is commonly found in yeasts and mammals and their involvement lies in vesicle fusion. The presence of vesicles among microsporidia is not a common phenomenon. Intracellular sorting or transport of cargo proteins from the atypical microsporidian golgi complex occurs by a mechanism that does not involve the participation of vesicles but rather tubular networks **(Beznoussenko *et al.,* 2007)**. Avesicular transport has been established through RT-PCR studies within one genus of microsporidia, *Paranosema,* classified as *Nosema* earlier, which infects grasshoppers, crickets and locusts **(Dolgikh *et al.,* 2010)**. Sequence similarity search showed the presence of more than hundred similar proteins within the UniProtKB database. These transporters have some important role to play in the parasite’s life cycle because their expression has been proven in spore and consecutive intracellular stages; however that role is unclear at present.

Four transporter families were common in at least two *Nosema* species, namely, Non-selective cation channel-2 (NSCC2) family (1.A.15), Major Intrinsic Protein (MIP) family (1.A.8), CorA metal ion transporter (MIT) family (1.A.35) and Nuclear pore complex (NPC) (1.1.1). NSCC2 proteins are homologues of general protein secretory pathway in yeast microsomes and have been found to act as non-selective cation channels in mammalian cytoplasmic membranes **(Tyerman, 2002)**. MIP family transporters are aquaporins in native form, but are also capable of transport of carbohydrates, glycerol, urea, ion etc. by an energy-independent mechanism **(Bienert *et al.,* 2008)**. Both these α-type channels have been reported in other microsporidia like *Encephalitozoon cuniculi, Enterocytozoon bieneusi* and *Anncaliia algerae,* and hence, can be considered to play similar roles in *Nosema sps* **(Peyretaillade *et al.,* 2012)**. Again, CorA transporters are similar to MscS transporters in substrate specificity, i.e, metal ions and has been reported previously in *E. bieneusi* **(Peyretaillade *et al.,* 2012)**. NPC homologues involved in transport across nuclear membrane were observed in *N. bombycis* and *N. antheraea* and not in other two microsporidia. Their presence has been also been reported in developmental stages of microsporidia previously **(Liu, 1972)**. These channels are conserved in eukaryotes, comprise of a number of nucleoporins and are responsible for nuclear cytoplasmic exchange. Apart from these, three species-specific transporter families (present only in one of the microsporidia) were discovered within Class 1- The autotransporter-1 (AT-1) family in *N. bombycis*; Hly III Family (1.C.113) and Rz/Rz1 Spanin1 Family (1.M.1) in *N. antheraea.* Both AT-1 and Rz(1) is involved in periplasmic transport of protein and hydrocarbons, respectively. The homologue of Hly family is a putative Hemolysin III-like protein which is currently uncharacterized. As evident from **Figure 1**, the presence of channels within the four species is quite variable. The ones infecting honeybees have less channels than the ones infecting silkworms. This uneven distribution is arising from the presence of Hly, Rz(1) and AT-1 families as well as the larger subset of homologues of NPC and MscS families in *N. antheraea* and *N. bombycis.* The presence of these additional channels in *N. bombycis* and *N. antheraea* poses an intriguing question as to what advantage does these extra transporters provide them regarding their host, the silkworms.

### [3.2] Class 2: Secondary carrier-type facilitators

Secondary carrier-type facilitators, also known as electrochemical-potential driven transporters, represent Class-2 of TCDB and employ a carrier-mediated process involving uniporters, symporters and antiporters for transport. We observed a total number of 101 transporters of *Nosema* belonging to Class 2 **(Table 3)**. *N. bombycis* had the highest number of secondary transporters (forty-two) out of all the four species studied here. All the *Nosema* transporters from Class 2 were classified as porters (uniporters, symporters and antiporters) and other families like ion-gradient driven energizers or transcompartment lipid carriers were absent.

**Table 3.**
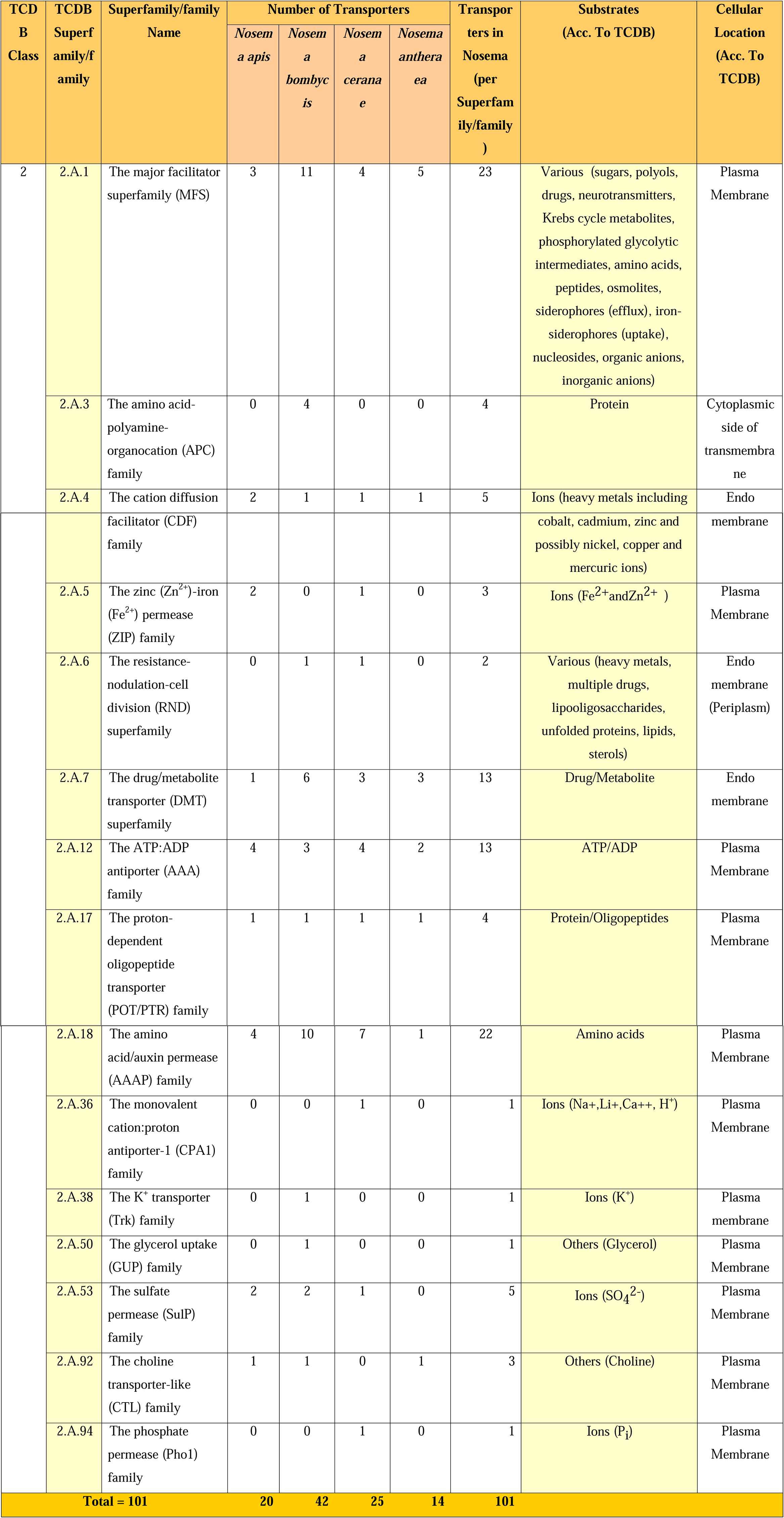
Distribution of Class 2 transporters with their substrates for *Nosema apis* (NA), *Nosema bombycis* (NB), *N. ceranae* and *N. antheraea:*

Out of the fifteen families, six were commonly found in all the four species, viz. Major facilitator superfamily (MFS) (2.A.1), Cation diffusion facilitator (CDF) superfamily (2.A.4), Drug/metabolite transporter (DMT) superfamily (2.A.7), ATP:ADP antiporter (AAA) family (2.A.12), Proton dependent oligopeptide transporter (POT/PTR) family (2.A.17) and Amino acid/auxin permease (AAAP) family (2.A.18). It is known that all microsporidian genomes encode two or more MFS transporters that have specificity for sugars imported from hosts **(Heinz *et al.,* 2014).** One of the MFS homologues belonging to sugar porter family (SP) (2.A.1.1) can take up environmental glucose to support its own metabolism. Numerous homologues of this family were present in *N. bombycis* (11 no.s). Other MFS families observed in *Nosema* were Proteobacterial intraphagosomal amino acid transporter (Pht) family (2.A.1.53), Unidentified Major Facilitator-14 (UMF14) Family (2.A.1.65) and the Drug-H+ antiporter (DHA1) family (2.A.1.2). These three families are commonly found in yeast and bacterial pathogens. The latter two have sequence similarities with multidrug-resistant transport proteins of yeast and *E. coli* as well as purine transporters **(Goffeau *et al.,* 1997)**. The second conserved Class 2 family is the Cation diffusion facilitator (CDF) family (2.A.4) responsible for heavy metal ion efflux and are mostly implicated in instances where human health and bioremediation is concerned **(Nies, 2003)**. It has also been reported to be involved in Co^2+^-ion uptake in *Saccharomyces cerevisae* **(Conklin *et al.,* 1992)**. *E. cuniculi* and *E. bieneusi* comprises of at least one transporter from this CDF family **(Peyretaillade *et al.,* 2012)**. Again, DMT superfamily plays a role in host adaptation and is associated with transport of variety of metabolites (nucleotide, sugar etc) from host cytoplasm **(Jack *et al.,* 2001).** Out of the 31 members of this superfamily, four were found within *Nosema,* viz., The Triose-phosphate Transporter (TPT) Family (acts as antiporter, exchanges organic phosphate ester for inorganic phosphate), The UDP-Galactose:UMP Antiporter (UGA) Family, The UDP-N-Acetylglucosamine:UMP Antiporter (UAA) Family (both act as antiporter, exchanges a nucleotide-sugar for nucleotide) and The NIPA Mg2+ Uptake Permease (NIPA) Family (mediates Mg^2+^ intake, causes a neurodegenerative disorder called Hereditary Spastic Paraplegia in humans and has homologues in other eukaryotes). The fourth common transporter is the AAA family, which has been reported in other microsporidia as well as bacteria, fungi and plants **(Tjaden *et al.,* 1999; Peyretaillade *et al.,* 2012)**. It is known that microsporidia lacks an ideal mitochondrion like structure and consequently, lacks the electron transport chain. Instead *Nosema* genus has acquired AAA transporters which can act as a part of the compensatory mechanism for the lack of the same and possibly have a role in importing ATP from host cytoplasm. AAAP family, another important transporter family, which is conserved across the *Nosema* genus, are associated with vacuolar bidirectional symport or antiport of various amino acids as well as ions (H+, Na+ etc) in lower eukaryotes such as yeast **(Russnak *et al.,* 2001; Chardwiriyapreecha *et al.,* 2010).** Finally, the proton-dependent oligopeptide transporter (POT/PTR) family involved in substrate (small peptides, oligopeptides, proteins etc.) efflux coupled to H^+^ antiport.

Apart from these conserved transporter families, four other were found in at least two of the *Nosema* species. Two of these families transport different types of ions, namely, the Zinc (Zn^2+^)-Iron (^Fe2+^) permease (ZIP) family (2.A.5) involved in acquisition of metal ion, especially in intake and maintenance of homeostasis of Zn^2+^; and the Sulfate permease (SulP) family which is involved in inorganic anion uptake. Other than these ion transporters, Resistance-nodulation-cell division (RND) superfamily (2.A.6), which is homologous to solute carrier family of mammals and can transport a range of substrates like peptides, amino acids (Histidine), nitrates etc. and the choline transporter-like (CTL) family (2.A.92), a solute carrier family for choline, were also identified.

A range of Class 2 transporters were also found to be species-specific. *N. bombycis* transportome had four specific transporter families- Amino acid-polyamine-organocation (APC) family (2.A.3) (functions as arginine/ornithine or cystine/glutamate antiporter to maintain cellular redox balance as well as cysteine/glutathione levels), Monovalent cation:proton antiporter-1 (CPA1) family (2.A.36) involved in Na^+^:H^+^ exchange, K^+^ transporter (TrK) family (2.A.38) (K^+^:H^+^ symport) and Glycerol uptake (GUP) family (2.A.50) (implicated in glycerol and amino acid uptake in bacteria, yeast etc.) **(Saier, 2000)**. Similarly, *N. ceranae* had only one unique transporter family, the phosphate permease (Pho1) family (2.A.94) which carries out inorganic phosphate transport in plants.

The number of transporters in *N. bombycis* is nearly twice to that in other three species, contributed by high number of MFS, DMT and AAAP family homologues. These three families are conserved across the four species and other microsporidia. A possible explanation of these high numbers of transporter homologues is that the rates of substrate exchange are higher in *N. bombycis.* The real question, however, is why this rate higher in *N. bombycis* than other three species.

### [3.3] Class 3: Primary active transporters

Primary active transporters of Class 3 drive transport of a solute against a concentration gradient using a primary source of energy. Around 32-37 primary transporters have been found to be present in other microsporidia (*E. cuniculi, E. bieneusi, A. algerae*) **(Peyretaillade *et al.,* 2012).** We identified eighty-three primary transporters in our subset of *Nosema* genus **(Table-4)**. All of these homologues drive the active uptake and/or extrusion of a solute or solutes via hydrolysis of diphosphate bond of inorganic pyrophosphate, ATP, or nucleoside triphosphate. Oxidation-reduction, methylation or decarboxylation driven transporters were not found in the studied *Nosema* genus.

**Table 4.**
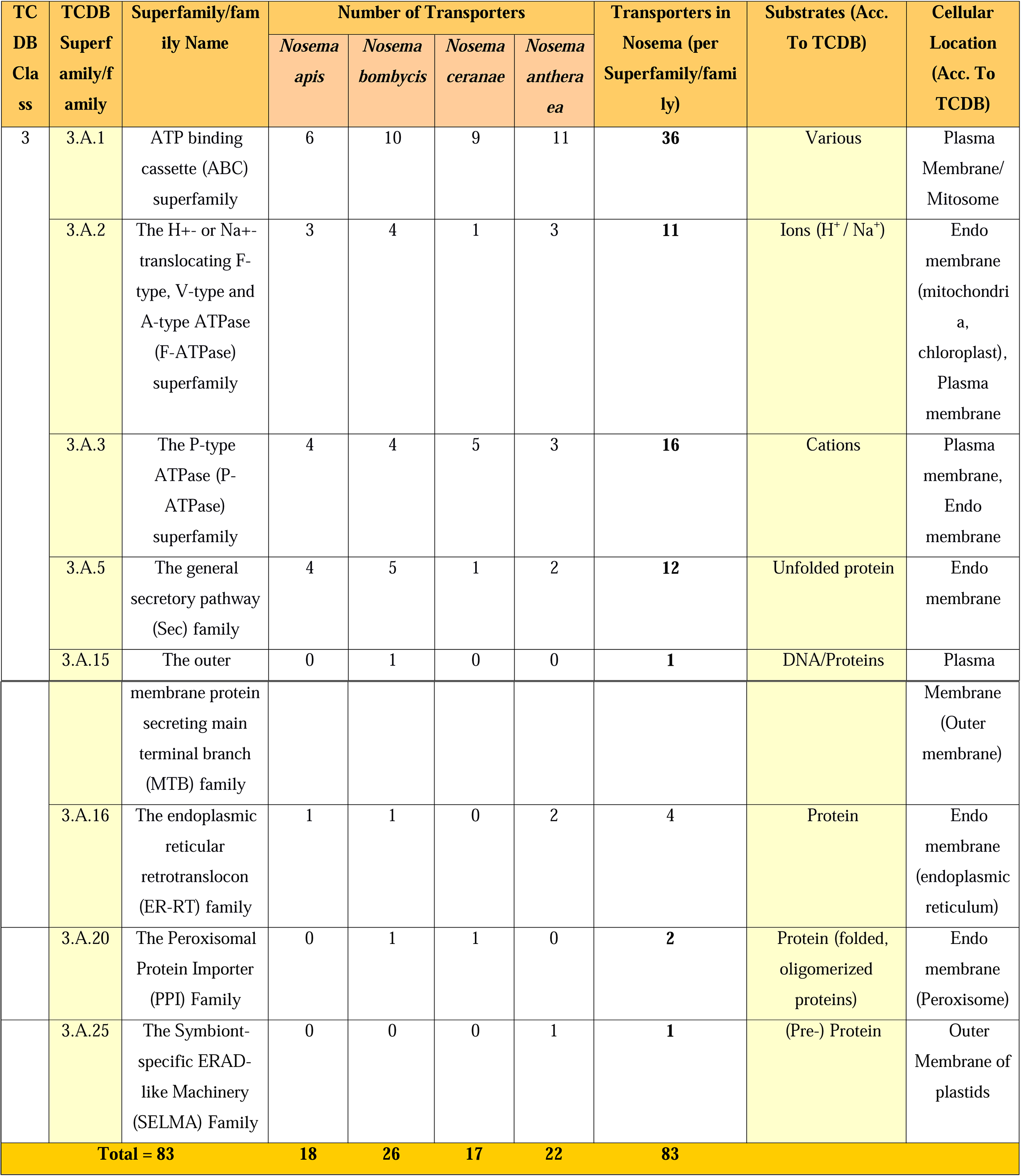
Distribution of Class 3 transporters with their substrates for *Nosema apis* (NA), *Nosema bombycis* (NB), *N. ceranae* and *N. antheraea*.

As per our observation, the largest group of Class 3 transporters were the ATP-binding cassette (ABC) transporters (3.A.1) superfamily, expressed uniformly across our studied *Nosema* subset. Similar observation was previously expressed in other microsporidia from *Encephalitozoon, Enterocytozoon* and *Anncaliia* genera **(Peyretaillade *et al.,* 2012)**. ABC transporters are associated with transport of various substrates like peptides, lipid, ions, drugs etc. A total of thirty-six ABC transporters have been identified in *Nosema,* whereas previous studies showed the presence of four sub-families in *E. cuniculi* **(Cornillot *et al.,* 2002)**. Within the ABC superfamily, Heavy metal transporter (HMT) family (also known as ABCB) (3.A.1.210) was the most abundant (21 no.s in the four *Nosema* sp.s), followed by Eye pigment precursor transporter (EPP) family (also known as ABCG) (3.A.1.204) (10 no.s in the four *Nosema sp.s).* Heavy metal transporters have been observed in *E. cuniculi* and *E. intestinalis* previously **(Cornillot *et al.,* 2002)**. Similarity between these putative transporters and that of yeast ATM1 protein which is a prototype for this subfamily suggests that they might be carrying out similar function in Fe–S cluster export. A related hypothesis proposes the role of cryptic mitochondria of microsporidia in iron-sulfur biogenesis **(Corradi, 2015)**. Similarly, EPP family of transporters have been identified previously in *E. cuniculi;* they are thought to be involved in guanine and tryptophan transport in *Drosophila melanogaster* **(Ewart *et al.,* 1994)**. Other than ABC superfamily, the classes 3.A.2 (The H+- or Na+-translocating F-type, V-type and A-type ATPase (F-ATPase) superfamily), 3.A.3 (The P-type ATPase (P-ATPase) superfamily) and 3.A.5 (The general secretory pathway (Sec) family) were conserved across the four *Nosema* species. F-ATPase superfamily is conserved from bacteria to eukaryotes and is associated with vacuolar transport of H^+^ and Na^+^. Only V-type ATPases were identified in *Nosema* genus. P-ATPases identified in this study were the ones involved in phospholipids translocation and cation transportation **(Alder-Baerens *et al.,* 2006)**. Again, homologues to yeast Sec-SRP complex (3.A.5.8) and mammalian Sec-SRP complex were observed in the fourth conserved and abundant transporter family in Class 3, i.e., Sec family, including Sec61 (α and γ) subunits and sec63 translocase subunits. Sec61 complex is the general secretory (Sec) pathway for protein secretion into ER with its subunits Sec61α, Sec61β, and Sec61γ. This complex has been identified previously in *N. bombycis, E. cuniculi* and *Antonospora locustae* **(Wu *et al., 2007*)**.

Two other families from Class 3 were expressed in at least two of the four organisms studied here, namely, Endoplasmic reticular retrotranslocon (ER-RT) family (3.A.16) and the Peroxisomal Protein Importer (PPI) Family (3.A.20). ER-RT transport proteins are abundant in *N. antheraea* and are supposedly associated with ER-associated degradation system which involves translocation of misfolded proteins from ER to cytoplasm for degradation. Microsporidia is known to lack any organelle like peroxisomes and the role of PPI transporters (3.A.20) needs in-depth analysis to understand the functions and relevance to its survival or other functions. Two species-specific unique transporter families from Class 3 were also found in *N. bombycis* and *N. antheraea,* indicating that it still retains several transporter genes which are absent in its *Nosema* counterparts. The outer membrane protein secreting main terminal branch (MTB) family (3.A.15) which is present only in *N. bombycis* and it shares 71% sequence identity with the pulF protein of *Klebsiella pneumoniae.* Presence of *K. pneumonia* in silkworm gut has been reported previously; the presence of this unique transporter family with bacterial-origin in *N. bombycis* is a probable case of horizontal gene transfer due to co-localization at a shared niche. The acquisition of this transporter could probably aid *N. bombycis* in DNA and protein transport. Another species-specific transporter found only in *N. antheraea* was the Symbiont-specific ERAD-like Machinery (SELMA) family (3.A.25) transporter, involved in transport of nucleus-encoded pre-protein in human pathogens, *Plasmodium falciparum* and *Toxoplasma gondii.*

## [4] Conserved and Unique transporters in *Nosema:* how are they relevant?

Microsporidia, being an obligate parasite with highly reduced gene content, is incapable of carrying out numerous processes that free-living organisms like yeast, bacteria etc. can carry out. These include Kreb’s cycle, electron transport chain, nucleotide synthesis, etc. With genome reduction via gene loss, the transportome diversity of a microsporidia should ideally decrease. Despite this, a set of core transporters like permeases allowing passage of sugar, glucose, metal ions viz. Mg^2+^, Ca^2+^; polyamine transporters; ion transporters with metal ion specificity; sulfate transporters etc. are commonly retained by any microsporidia **(Weiss and Becnel, 2014)**. *Nosema,* like any other microsporidia, is also dependent on the host cytoplasmic contents for survival and proliferation. To obtain a plausible view of the core transporters of *Nosema* genus, we compared the transportome of *N. apis, N. bombycis, N. ceranae* and *N. antheraea* **(Figure 3)**. Then, we compared the existing data on reviewed and unreviewed microsporidian transporters from UniProt to the complete set of *Nosema* transporters.

**Figure 3.**
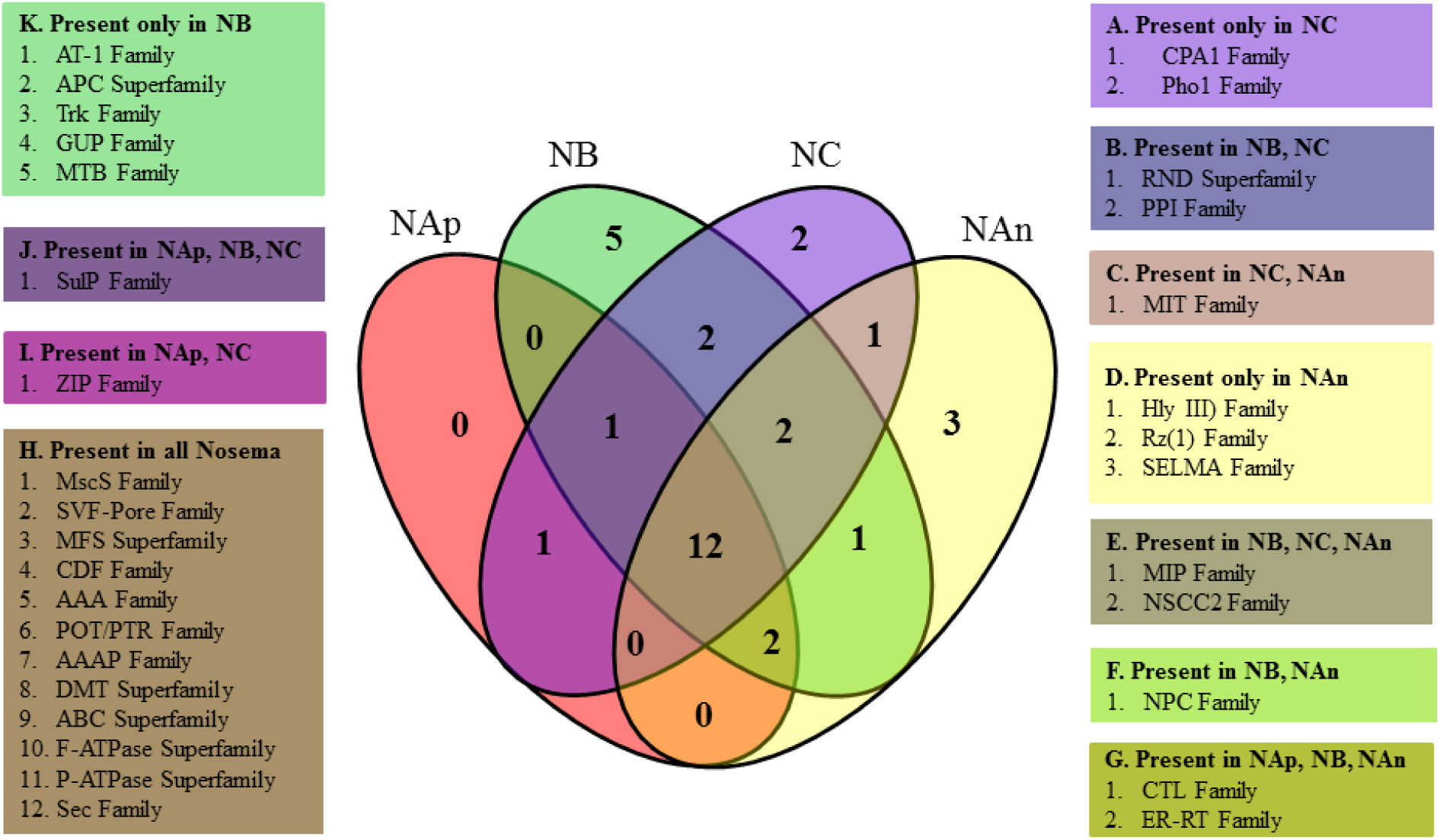
Transporter family distribution among the *Nosema* genus: Venn diagram showing shared and unique transporter families among four *Nosema* species, viz. NAp- *Nosema apis,* NB- *N. bombycis,* NC- *N. ceranae,* NAn- *N. antheraea*.

Our study revealed that the *Nosema* genus too, has a core set of transporter families conserved among them **(Figure 3 and 4)**. This set of twelve transporter families (ABC, DMT and MFS superfamily, F-ATPase, P-ATPase, Sec, MscS, CDF, SVF-Pore, POT/PTR, AAA and AAAP family) can be perceived to be crucial for a typical microsporidian life **(Figure 4)**. Except the SVF-Pore family, all others have been reported in at least one microsporidian species other than those in *Nosema* genus **(Refer to S-3)**. Three of these families are involved in ion transport, namely, the energy-independent MscS family and the energy-dependent F-ATPase and P-ATPase families. Recent research on microsporidia like *T. hominis, N. parisii* and *E. cuniculi* has reported that microsporidian MscS have two distinct origins, one is eukaryotic MscS1 proteins and another is bacterial-like microsporidian MscS2 **(Heinz *et al.,* 2012; Cuomo *et al.,* 2012; Grisdale *et al.,* 2013)**. MscS proteins can regulate osmotic homeostasis at the cell surface during different life stages by opening or closing a channel permeable to water and small ions in response to mechanical deformation of the cell membrane, such as that caused by physical or osmotic pressure **(Nakjang *et al.,* 2013)**. It is to be noted that increase or decrease osmotic pressure in an intracellular parasite can be related to increase or decrease in membrane tension to speed up or delay its egress from a host cell **(Lavine and Arrizabalaga, 2008)**. This means that MscS family could have a probable role in *Nosema* infection process. Again, the V-type ATPase (from F-ATPase family) and the P-ATPases mediate ATP-dependent H+/Na ion transport and have been widely observed in other microsporidian species across different genera such as *Anncaliia, Encephalitozoon, Enterocytozoon* etc **(Peyretaillade *et al.,* 2012)**. Both these families are capable of cation efflux and can be speculated to be related to pH regulation, i.e, maintenance of resting pH and recovery from pH dysregulation inside the gut of a host organism by H+ extrudation. This phenomenon has been experimentally observed previously in protozoa such as *Plasmodium falciparum* **(Saliba and Kirk, 1999)**. Similar to *P. falciparum,* the hosts for *Nosema* genus are also insects and microsporidian spore germination occurs in various parts of the midgut or gut epithelial cells in insects of Hymenoptera and Lepidoptera **(Weiss and Becnel, 2014)**. Presence of these transporters probably helps *Nosema* sp.s in cation uptake, not only for nutritional or metabolic purposes but also survival amidst its hosts. Similar roles may also be played by the other core ion transporter family, CDF and as well as less conserved SulP, NSCC2 etc.

**Figure 4.**
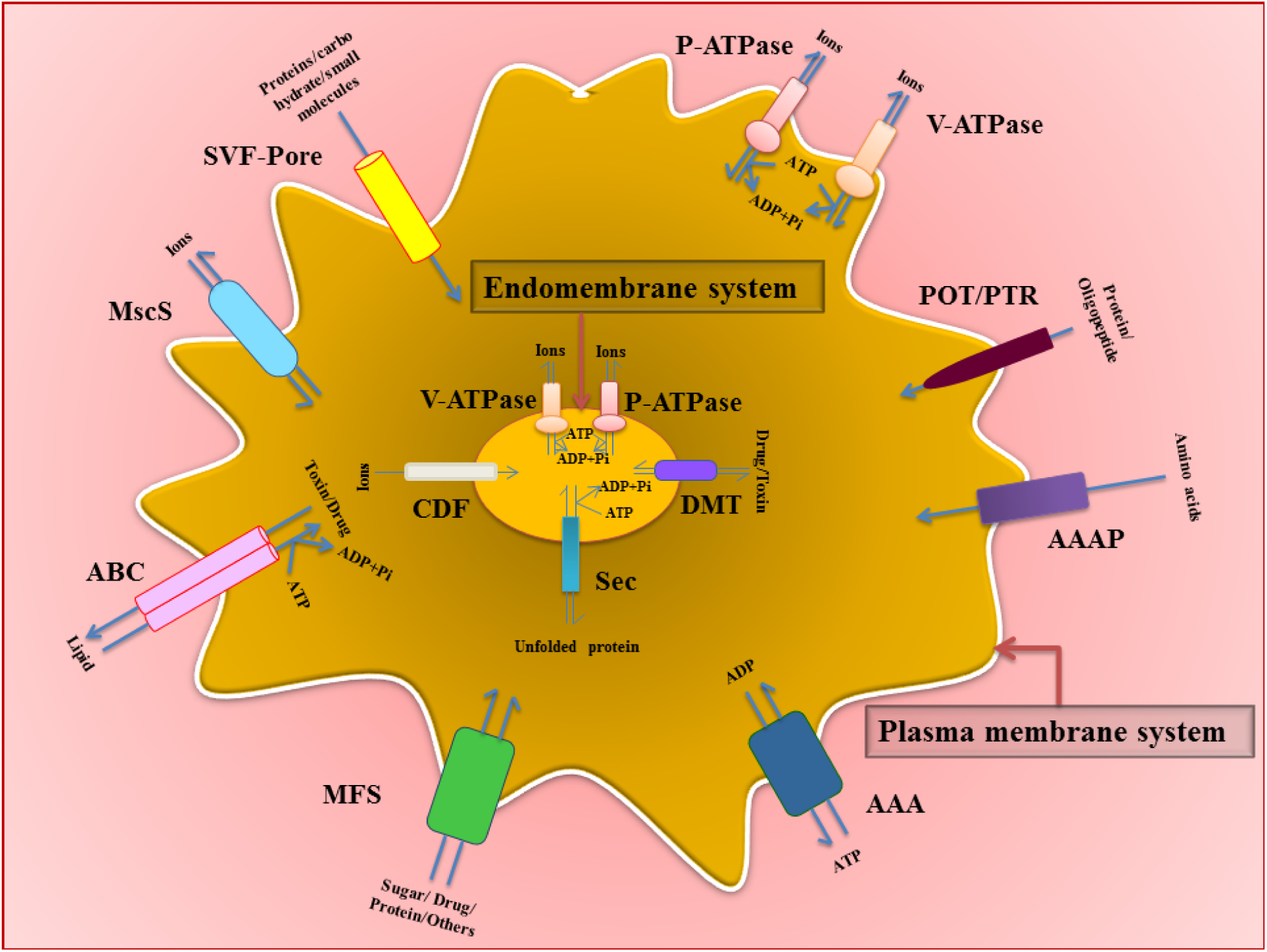
Diagrammatic representation of a typical *Nosema* cell with the core set of transporters conserved within the *Nosema* genus. *(Note- The images used to denote the various type of transporter families doesn’t imply their acutal stucture within plasma or endomembrane)*

The MFS superfamily is one of the predominant transporter families found in microsporidia and study of the *Nosema* genus provided us a similar observation. All known microsporidian genomes encode three or more MFS transporters with specificity for sugars likely to be acquired from hosts **(Heinz *et al.,* 2012)**. Microsporidian genome retains the pathways like glycolysis, pentose phosphate pathway, etc. but expression of the same varies from species to species and also from the habitat nature of the parasite (aquatic/terrestrial) **(Undeen and Vander Meer, 1999)**. In two of the *Nosema* species (*N. apis* and *N. ceranae*) studied here, it has been observed that sugar utilization increases along with spore counts within the host, resulting in energetic stress to the host that translates into several abnormal behavioral manifestations **(Martín-Hernández *et al.,* 2011)**. These studies, combined with our results portray the possibility that pathways for sugar uptake and utilization for ATP synthesis could still be functional within the *Nosema* genus (except *N. ceranae* which lacks this gene according to genome analysis). Presence of a terminal electron acceptor enzyme found in mitosomes, Alternative oxidase, has been hypothesized to be present in some microsporidia, aiding in glycolytic pathway of energy generation **(Williams *et al.,* 2010)**. Other than sugar, MFS family also transports other substrates like polyols, amino acids, osmolites, drugs, neurotransmitters, Krebs cycle metabolites, phosphorylated glycolytic intermediates, inorganic anions, etc. making it an inevitable part of the *Nosema* as well other microsporidian life cycle. Another substrate importing transporter family which was less conserved among the studied genus was the MIP family.

Among the highly conserved families were the Sec, POT/PTR, SVF-Pore families involved in polypeptide/protein transport. Homologues of the Sec family have been observed in microsporidia previously **(Peyretaillade *et al.,* 2012)**. It was hypothesized that microsporidia exploits a protein transport and secretion machinery which is similar to yeast or mammalian cells **(Weiss and Becnel, 2014)**. However, even this machinery has undergone severe loss of nonessential genes. *E. cuniculi* genome reportedly, has all the components of the Sec61 translocon channel along with other associated proteins like Sec62-63, Hsp70 etc. which make translocation of proteins into ER possible **(Katinka *et al.,* 2001)**. Our analysis discovered homologues of Sec61 (α and γ) subunits and sec63 translocase subunits and high conserved nature of these homologues across the four species of *Nosema* genera suggests that this pathway might be functional in *Nosema,* carrying out co-translational translocation of polypeptide chains into the ER lumen. Another complimentary mechanism serving this purpose is the SNARE protein mediated vesicular fusion. Homologues of the SVF-Pore family (1.F.1.1) that consists of proteins from the SNARE complex was conserved across *Nosema* (during our study) and *E. cuniculi* (from the UniProt database). Apart from the functions mentioned above, these transporter families might also be involved in other forms of intracellular transport and exocytosis.

Multiple studies have suggested the *de novo* synthesis pathway for amino acids is lost in microsporidians, making them partially dependent on host-derived amino acids and partially on self-mediated inter-conversion **(Heinz *et al.,* 2012)**. In such a case, transporters with differential specificity for differently charged amino acids like the ones in AAAP family can be of immense importance for the organism. Apart from the functionalities of this family discussed in Section 3.2, these core transporters also have a role in the pathogenicity of *Nosema.* Usage of host-derived amino acids disrupts its levels within the host, indirectly putting it in energetic stress. The life cycle of *Nosema* demands usage of considerable amounts of amino acids, for eg.- during spore wall or polar tube formation. Whether the levels of amino acid depletion in host are relative to the rate of infection by *Nosema* can be experimentally determined, as has been done in case of sugar usage. This will provide another measure of energetic stress that this intracellular parasite places on its insect hosts.

As for the most important energy component of any organism, i.e., ATP, microsporidia is dependent on hosts (although presence of an alternative glycolytic pathway has been hinted previously) and the conserved AAA family is capable of serving this purpose for *Nosema* and its microsporidian counterparts. Regarded as a genomic hallmark for microsporidia, these transporters have similarities to bacteria and are speculated to have been acquired by horizontal gene transfer led by co-existence. As discussed in section 3.2, AAA proteins can exchange ATP for ADP, gaining energy in form of an extra pyrophosphate bond. These ATP exchanges along with that of sugar and amino acids further distresses host metabolism. So, these transporters have been used a targets of gene silencing to reduce parasite load in Nosemosis, leading to favorable changes in host physiology **(Paldi *et al.,* 2010)**. The putative transporters discovered in this study can be further characterized and experimentally tested as novel gene silencing targets. Again, establishing the importance of DMT superfamily within *Nosema* genus and microsporidia, as a whole, can be related to the myriad of substrates it deals with. For example-Mg^2+^ ion transported by NIPA family can be utilized by *Nosema* augmenting the adherence of spore wall to host cell surface as observed in *E. cuniculi* glycosaminoglycans **(Southern *et al.,* 2006)**. Endosomal transporter families were also found amongst other members of DMT superfamily, including those for golgi apparatus. However, these organelles are highly reduced or absent in microsporidia and more evidence on their presence is required before arriving at any conclusion. Finally, the ABC superfamily, discussed in Section 3.3, is found in many other microsporidia and the present study concurs with this fact. The presence of this family in the microsporidian genomes, like *E. cuniculi,* have been confirmed but with unknown specificity **(Cornillot *et al.,* 2002)**. ABC transporters, reportedly act as exporters in certain parasitic protists (the classification that *Nosema* previously belonged to) but they might have a role to play in import or retrieval of host nutrients as well; its presence of *Nosema* and other microsporidia indicates the same. Its involvement in unidirectional transportation of substrates including extrusion of anti-parasitic molecules within eukaryotic parasites as a measure of self-defense is a possibility.

From the standpoint of an individual species, it *N. bombycis* appears to be the most endowed or rather, most progressive microsporidian from *Nosema* genus. With an arsenal of 78 transporter proteins, it should have an advantage over the other three species studied here in terms of survival, pathogenicity and proliferation. This might be the reason behind the broad range of domesticated and wild Lepidopteran hosts that it can infect; susceptible insect families include Bombycidae, Noctuidae, Pieridae, Arctiidae and Crambidae **(Kashkarova and Khakhanov;Kudo and DeCoursey, 1940).** *N. bombycis* transportome also constitutes of the largest subset of unique, species-specific transporters, which could be result of adaptive evolution or horizontal gene acquisition from co-existing gut microbes. *N. bombycis* is closely followed by *N. antheraea* (51 no.s) in terms of transporter count. The host of *N. antheraea* is a wild silkworm species, *A. pernyi,* unlike the hosts of *N. apis* and *N. ceranae* species studied here **[Table-1].** Like *N. bombycis,* this organism also has three unique transporter families which have not been reported in other microsporidians till now. How these unique proteins aid this obligate parasite is an intriguing question.

Observation of the transporters according to host specificity shows that the share of ion transporters differs between the honey bee and silkworm parasites **(Refer Table 2**, **3** and **4)**. Out of the four observed ion transporter families in *N. apis,* three are conserved while one is common with *N. ceranae* only. However, unique ion-transporter families discovered in *N. bombycis* points out that it may involve these transporters in pH or osmotic stress regulation, thus conferring it some added advantage in its survival within the cytoplasm of an array of insect hosts. The absence of same in *N. apis,* supposedly, confers a disadvantage in its host adaptation abilities.

Close observation of transporter distribution across the four species shows a pattern where the *N. apis, N. bombycis and N. ceranae* has less number of porters/channels (Class 1) and more number of secondary carriers (Class 2) **(Figure 2)**. However, *N. antheraea* transportome shows a reverse pattern. Within the core set of ten transporter families in *N. antheraea,* number of transporter proteins is reduced in AAA, AAAP, MFS etc. Again, the number of homologues of NPC family within *N. antheraea* is high, increasing the share of channels/porters within its transportome. It is somehow related to the fact that *Nosema* uses host nucleus as its developmental niche **(Corradi, 2015)**. It is also noteworthy that within *N. antheraea* transportome, the number of primary transporters is highest (22 no.s). How does a highly host-dependent, energy-efficient intracellular parasite like *N. antheraea* keep up with the energetic costs for Class 3 when the number of ATP transporter is as low as two? **(Table-3)** Assuming that all the deduced transporters of Class 3 are functional, a striking hypothesis can be made from the above observations- *N. antheraea* have compensated for the dearth of Class 2 transporters by scaling-up the transporters from Class 1 and 3. From **Table-1**, it is apparent that *N. antheraea* has the smallest genome among four organisms and among other genes, it may have shed the genes for secondary carrier facilitators. In terms of species-specific transporters, *N. antheraea* and *N. bombycis* are richer than *N. apis* and *N. ceranae.* However, the ones exclusively found in *N. antheraea* like Rz(1) or Hly III have not been reported earlier and requires further analysis. Our observations are largely based on the outputs of an *in silico* pipeline and requires greater in-depth experiments before garnering any conclusive view. A total of 10 unique species-specific transporter families are present in the *Nosema* genus **[Figure 4]**. The importance of these transporters in the microsporidian should be experimentally verified.

## [5] Conclusion

The current scenario of apiculture and sericulture is threatened by microsporidians of *Nosema* genus and new strategies are required to tackle these parasites. Since microsporidia are entirely dependent for some crucial substrates like ATP, sugar, nucleotides etc. on its host, the transporters for these substrates can act as critical components of a pest management strategy. The broad spectrum of transporter proteins within the *Nosema* genus identified by us using available data can act as a valuable resource for future studies. This includes understanding microsporidian biology, inner mechanism and its relation to host variability.

## [6] Acknowledgement

UB thanks the Department of Biotechnology, Govt. of India, New Delhi for supporting the research through the UXCEL project (Sanction Order No: BT/411/NE/U-Excel/2013 dated 06.02.2014) and Institutional Biotech Hub (Project BT/04/NE/2009), for providing the necessary facilities to carry out the study. HC, DK and SS express gratitude towards MHRD and IITG for financial support in the form of fellowship.

## [9] List of Supplementary Data

### Supplementary Data 1 (S-1)

List of putative transporter proteins with UniProt/SilkPathDB ID and the following corresponding data-

- UniProt/SilkPathDB ID
- TCDB ID
- TCDB Family/Superfamily Name
- Number of predicted transmembrane domains according to HMM**-**TOP, TMHMM and PHOBIUS

### Supplementary Data 2 (S-2)

List of TCDB families conserved and unique in *Nosema* genus

### Supplementary Data 3 (S-3)

List of characterized and putative transporter families in microsporidian species according to UniProt (As on March, 2016) and their comparison to total transportome of *Nosema.*

